# Surface LSP1 is a phenotypic marker distinguishing human classical versus monocyte-derived dendritic cells

**DOI:** 10.1101/813410

**Authors:** Sandrine Moutel, Anne Beugnet, Aurélie Schneider, Bérangère Lombard, Damarys Loew, Sebastian Amigorena, Franck Perez, Elodie Segura

## Abstract

Human mononuclear phagocytes comprise several subsets of dendritic cells (DCs), monocytes and macrophages. Distinguishing one population from another is challenging, especially in inflammed tissues, due to the promiscuous expression of phenotypic markers. Using a synthetic library of humanized llama single domain antibodies, we identified a novel surface marker for human naturally-occuring monocyte-derived DCs. Our antibody targets an extra-cellular domain of LSP-1, specifically on monocyte-derived DCs, but not on monocytes, macrophages or classical DCs. Using this antibody, we provide evidence that the recently described blood DC3 population does not correspond to circulating monocyte-derived DCs. Our findings will pave the way for a better characterization of human mononuclear phagocytes in pathological settings.

## Introduction

Mononuclear phagocytes play a central role in the initiation and resolution of innate and adaptive immune responses. They comprise several populations of monocytes, macrophages and dendritic cells (DCs), which are classically defined by their phenotype and ontogeny (1). Monocytes are very plastic cells, and can differentiate into cells displaying typical features of macrophages or DCs, referred to as monocyte-derived macrophages (mo-Mac) and monocyte-derived DCs (mo-DC). This phenomenon has been described *in vitro* and *in vivo*, in both mouse and human (2–4). Recently, high resolution single-cell technologies have revealed a new subset of blood DCs, termed DC3, displaying a mixed transcriptomic profile of classical DC (cDC) and monocyte genes (5–7), reminiscent of that of mo-DC (8, 9). Whether blood DC3 represent circulating mo-DC has been unclear.

Accumulating evidence suggests that monocyte-derived cells are involved in the pathogenesis of autoimmune and chronic inflammatory diseases (3). However, their characterization in inflammed tissues is complicated by the promiscuous expression of numerous phenotypic markers, shared in particular with macrophages and cDCs. More specific markers are needed in order to advance our understanding of the respective properties of macrophages, cDCs and mo-DC, and ultimately to allow the manipulation of these cells for therapeutic strategies.

To identify novel markers for human mo-DC, we have used a synthetic library of humanized llama single domain antibodies. We identified one antibody, recognizing surface LSP-1, that stains specifically mo-DC, but not monocytes, macrophages, cDCs or DC3.

## Results

In order to identify novel surface markers for human naturally-occuring mo-DC, we set up a phage display screen using a synthetic library of humanized llama single domain antibodies (termed VHH) (10). We have previously shown that peritoneal ascites from cancer patients contain a population of mo-DC (8, 11). Cells from tumor ascites were separated into DCs and all other cells (non-DCs), including tumor cells, macrophages, T cells and other immune cells. The library was first depleted for phages binding to non-DCs, then phages binding to ascites DCs were screened (fig 1A). For subsequent studies, we produced the VHH of interest with a human Fc region containing a Streptavidin-binding peptide. Using this strategy, we identified a VHH antibody, termed "D4", that stains ascites DCs but not macrophages from the same samples (fig 1B). We have recently reported a culture model allowing the generation of *in vitro* counterparts of naturally-occuring mo-DC and mo-Mac, by culturing CD14^+^ monocytes with M-CSF, IL-4 and TNFα (9). In this model, the VHH D4 stained mo-DC but not mo-Mac (fig 1C). We have also shown that the classical model of culturing monocytes with GM-CSF and IL-4 generates mo-DC that do not closely resemble the ones found *in vivo* in inflammatory fluids (9). Consistent with this, the VHH D4 did not stain mo-DC generated with GM-CSF and IL-4 (fig 1C).

**Figure 1.**
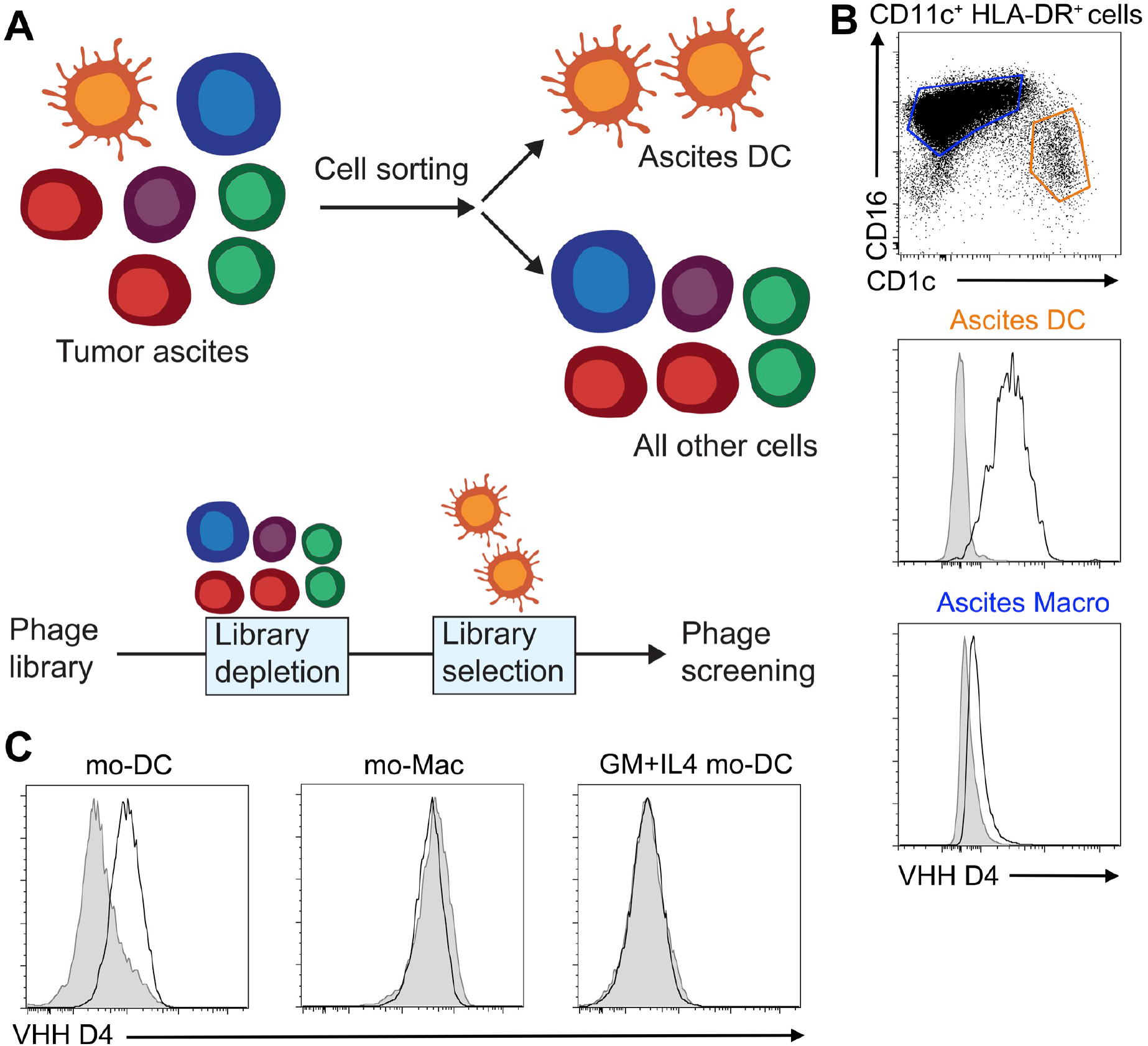
Identification of a VHH specific for monocyte-derived dendritic cells. (A) Strategy for identifying novel surface markers for ascites dendritic cells (DC). (B) Ascites cells were stained with VHH D4. Stainings for DC and macrophages (Macro) are shown (representative of 5 individual donors). Gray shaded histograms are fluorescence-minus-one controls. (C) Monocytes were cultured with M-CSF, IL-4 and TNFa to generate DC (mo-DC) and macrophages (mo-Mac), or with GM-CSF and IL-4 to generate DC (GM+IL4 mo-DC). Cells were stained with VHH D4. Gray shaded histograms are fluorescence-minus-one controls. Representative of 4 individual donors.

To determine the target of VHH D4 on the surface of mo-DC, we performed immunoprecipitation followed by proteomics mass spectrometry, using *in vitro*-generated mo-DC (cultured with M-CSF, IL-4 and TNFα) as a source of material. Mass spectrometry identified only one protein: Lymphocyte-Specific Protein 1 (LSP-1), a protein of predicted molecular weight of 60 kDa. To validate this result, we performed immunoprecipitation with VHH D4 on *in vitro*-generated mo-DC and mo-Mac, and revealed the immunoprecipitated proteins using Streptadivin (fig 2A) or a commercial anti-LSP-1 antibody (fig 2B). Both methods showed the same band around 55 kDa, only in immuno-precipitated material from mo-DC. This observation confirms that the VHH D4 binds LSP-1. LSP-1 is a F-actin binding protein reported to be expressed in all leukocytes (12). Using our previously generated transcriptomic data (9), we confirmed that *LSP1* was expressed at similar levels in ascites DC, ascites macrophages, *in vitro* mo-DC and mo-Mac generated with M-CSF, IL-4 and TNFα, *in vitro* mo-DC generated with GM-CSF and IL-4, and blood monocytes (fig 2C).

**Figure 2.**
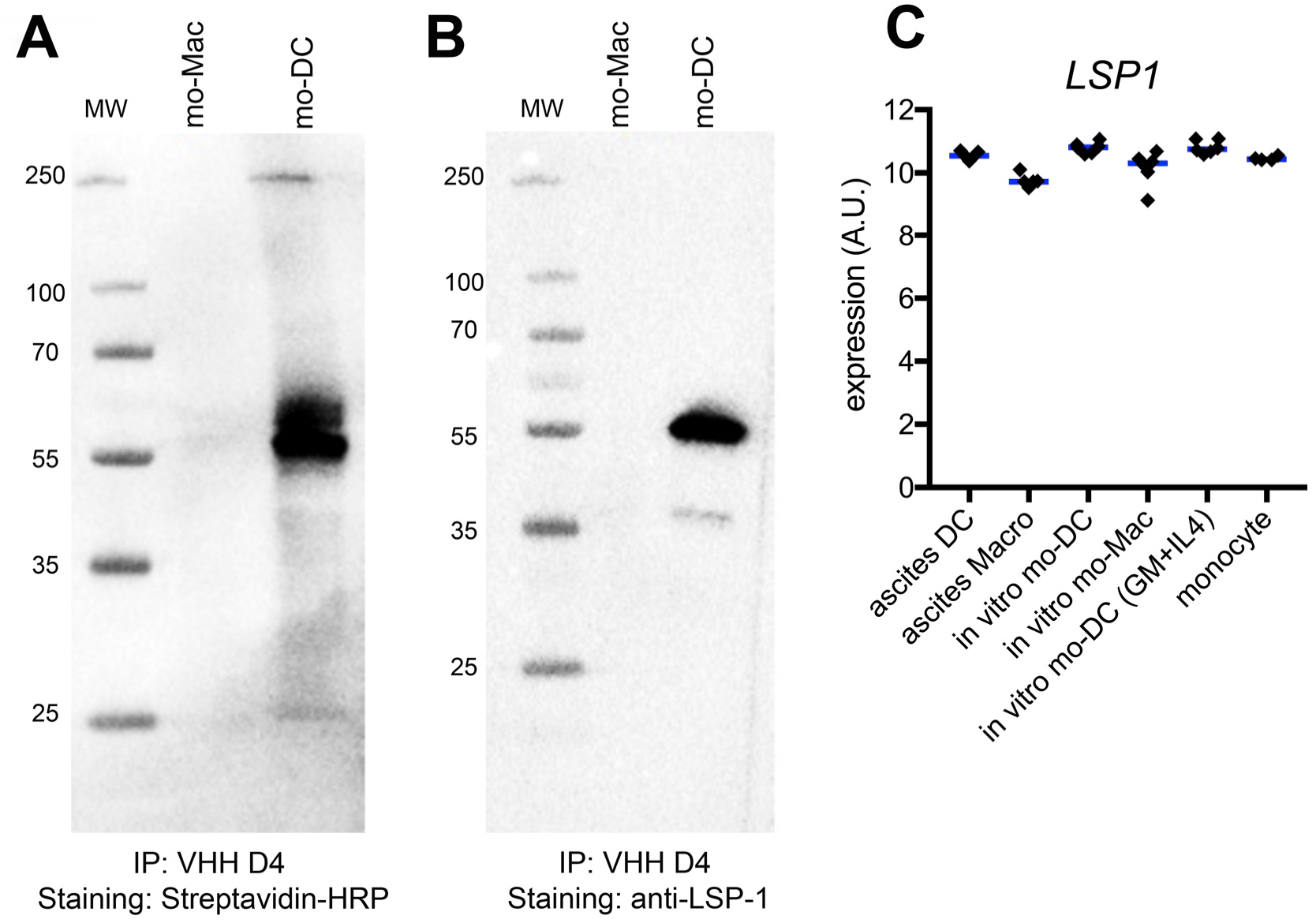
Identification of LSP-1 as the target of VHH D4. *In vitro*-generated mo-DC or mo-Mac were incubated with VHH D4, then lysed. Immuno-precipitation was performed on cell lysate using the streptavidin-binding peptide tag. MW = molecular weight. Immuno-precipitated material was analyzed by Western Blot. Staining was performed using streptavidin-HRP (A) or a commercial anti-LSP-1 antibody (B). (C) Expression levels (arbitrary units) of *LSP1* in ascites DC, ascites macrophages, *in vitro* mo-DC and mo-Mac generated with M-CSF, IL-4 and TNFα, *in vitro* mo-DC generated with GM-CSF and IL-4, and blood monocytes. Each dot represents an individual donor. Median is shown. Affymetrix data from dataset GSE102046.

We then sought to address whether the VHH D4 could also stain other myeloid cells, in particular cDCs. We first used the VHH D4 on cells isolated from human tonsils. We have previously shown using single-cell RNA-seq analysis that tonsils contain cDC1 (CD141^high^CD1c^−^ DCs) and cDC2 (CD141^low^CD1c^+^ DCs), but no population of mo-DC (13). There was no significant staining of VHH D4 on cDC1, cDC2, nor tonsil macrophages (fig 3A). This result suggests that the VHH D4 is specific of mo-DC, and does not stain cDCs.

**Figure 3.**
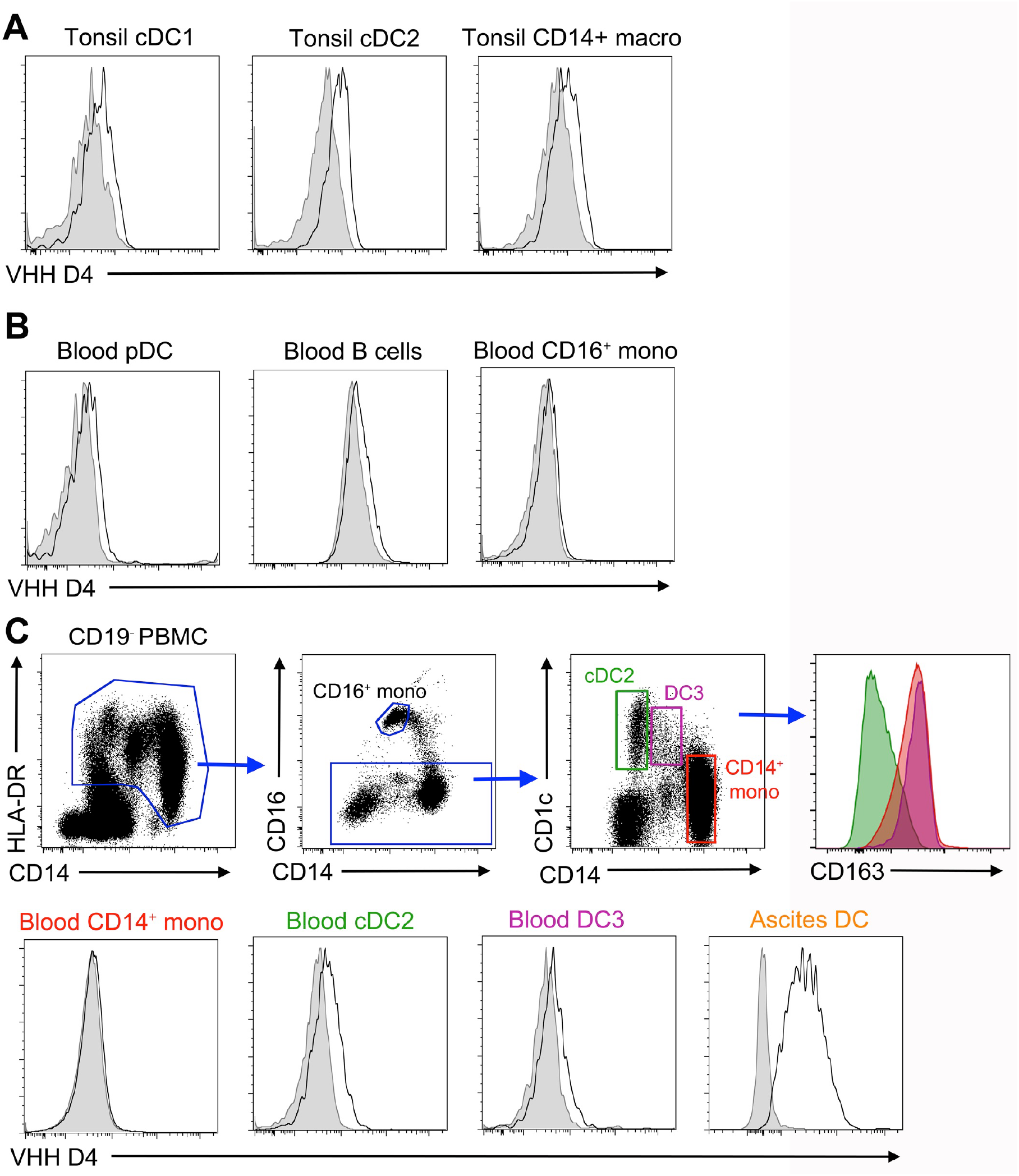
VHH D4 does not stain classical dendritic cell populations. (A) Tonsil DC and macrophages were stained with VHH D4. Gray shaded histograms are fluorescence-minus-one controls. Representative of 2 individual donors. Peripheral blood mononuclear cells (B-C) or ascites cells (C) were stained with VHH D4. Gray shaded histograms are fluorescence-minus-one controls. Representative of 4 (blood) or 4 (ascites) individual donors. (C) Gating strategy for analyzing CD14^+^ monocytes, cDC2 and DC3 is shown. Staining of ascites DC is shown as a positive control.

To extend these observations, we stained blood peripheral blood mononuclear cells. There was no significant staining on pDC, B cells nor CD16^+^ monocytes (fig 3B). It has recently been shown that blood CD1c^+^ DCs are heterogenous and comprise CD1c^+^CD14^−^CD163^−^ DCs (*bona fide* cDC2) and CD1c^+^CD14^+^CD163^+^ DCs, termed DC3 (5-7, 14). The VHH D4 did not show a significant staining on blood CD14^+^ monocytes, cDC2 nor DC3 (fig 3C), suggesting that DC3 are not blood-resident mo-DC.

## Discussion

Using a synthetic library of humanized llama single domain antibodies, we have identified one antibody, recognizing LSP-1, that is specific for human mo-DC and distinguishes them from monocytes, macrophages, cDCs or DC3.

LSP-1 has been shown so far to reside on the cytoplasmic side of the plasma membrane (15). After membrane permeabilisation, LSP-1 can be detected by flow cytometry, in particular in B cells and macrophages (12). Because our stainings with VHH D4 were performed without membrane permeabilisation, our results suggest the existence of an extra-cellular domain of LSP-1 specifically in mo-DC. The VHH D4 stains both mo-DC found *in vivo* in tumor ascites and generated *in vitro* with M-CSF, IL-4 and TNFα, indicating that the expression of a surface domain of LSP-1 is not due to the inflammatory micro-environment, but rather as the result of a mo-DC core transcriptional program. Whether surface LSP-1 results from alternative splicing or another post-transcriptional mechanism remains open for future investigation.

DC3 express a mixed transcriptional program with hallmark cDC2 genes (such as *FCER1A, CD1C, CLEC10A*) as well as typical monocyte genes (such as *S100A8, S100A9, VCAN*) (5–7), raising the question of their ontogeny, in particular whether DC3 correspond to circulating mo-DC. The VHH D4 did not show a significant staining on blood DC3, suggesting that they do not represent circulating blood mo-DC. This result is consistent with recent work showing the dependence of blood DC3 on Flt3-ligand (6), a characteristic feature of cDCs.

Together with our previous phenotypic characterization of mo-DC (9) and recent single-cell analysis of blood cDC2, DC3 and monocytes (5, 6), our work enables a more precise phenotypic definition of human DC subsets. cDC2 can be defined as CD88^−^CD14^−^CD1c^+^FcεRI^+^CD226^−^CD163^−^sLSP1^−^, DC3 as CD88^−^CD14^+^CD1c^+^FcεRI^+^CD226^−^ CD163^+^sLSP1^−^ and mo-DC as CD88^low^CD14^+^CD1c^+^FcεRI^+^CD226^+^CD163^−^sLSP1^+^.

Mo-DC have been identified in several inflammatory contexts in human tissues (3). In cancer, the picture is less clear as DCs displaying a phenotype consistent with both mo-DC and DC3 have been reported in colorectal, breast and lung tumors (16–18). DC3 are increased in the blood of patients with melanoma (7) and systemic lupus erythematosus (6), while blood DC3 infiltrate the lung alveolar space upon LPS-induced acute inflammation (19). The respective role of these two DC subtypes in cancer or inflammation remains elusive. Our findings will pave the way for a better characterization of DC3 and mo-DC in pathological settings, ultimately allowing their manipulation for therapeutic strategies.

## Methods

### Human Samples

Buffy coats from healthy donors (both male and female donors) were obtained from Etablissement Français du Sang (Paris, France) in accordance with INSERM ethical guidelines. Tumor ascites from ovarian cancer patients were obtained from Hôpital de l’Institut Curie in accordance with hospital guidelines. Tonsils from healthy patients (both male and female) undergoing tonsillectomy were obtained from Hôpital Necker (Paris, France). According to French Public Health Law (article L1121), written consent and IRB approval are not required for human non-interventional studies.

### Cell isolation

Tonsil samples were digested as described previously (20). In brief, samples were cut into small fragments, digested with 0.1 mg mL^−1^ Liberase TL (Roche) in the presence of 0.1 mg mL^−1^ DNAse (Roche) for 40 minutes at room temperature before addition of 10 mM EDTA. Cells were filtered on a 40 μm cell strainer (BD Falcon) and washed. Light density cells were isolated by centrifugation on a Ficoll gradient (Lymphoprep, Greiner Bio-One). DCs and macrophages were enriched by depletion of cells expressing CD3, CD15, CD19, CD56 and CD235a using antibody-coated magnetic beads (Miltenyi). Peripheral Blood Mononuclear Cells (PBMC) were prepared by centrifugation on a Ficoll gradient. Blood CD14^+^ monocytes were isolated from healthy donors’ PBMC by positive selection using anti-CD14-coated magnetic beads according to manufacturer’s instructions (Miltenyi). DCs and macrophage populations from ascites were isolated after cell sorting on a FACSAria instrument. Ascites DCs were gated as HLA-DR^+^CD11c^+^CD1c^+^CD16^−^.

### Cell culture

Blood monocytes (1×10^6^ cells mL^−1^) were cultured for 5 days in RPMI-Glutamax medium (Gibco) supplemented with antibiotics (penicillin and streptomicin) and 10% FCS in the presence of 100 ng mL^−1^ M-CSF (Miltenyi), 5 ng mL^−1^ IL-4 (Miltenyi) and 5 ng mL^−1^ TNF-α (Miltenyi), or 100 ng mL^−1^ GM-CSF (Miltenyi) and 5 ng mL^−1^ IL-4 (Miltenyi). In some experiments, mo-DC and mo-Mac were purified by cell sorting on a FACSAria instrument. Mo-DC were gated as CD1a^+^CD16^−^ and mo-mac as CD1a^−^CD16^+^.

### Phage display screening

A synthetic phage display library of humanized llama single domain antibodies (NaLi-H1 library) was used as described (10) to select binders. Screening of positive clones was performed using 100 µL of supernatant (80 µL phages + 20 µL PBS/human serum 1%) incubated for 1 h on ice with 1×10^5^ cells from tumor ascites. After washing, phage binding on ascites cells was detected by flow cytometry using an antibody against M13, and ascites DCs were gated as HLA-DR^+^CD11c^+^CD1c^+^CD16^−^ and ascites macrophages as HLA-DR^+^CD11c^+^CD1c^−^CD16^+^.

### Western blot

Cells were lysed in RIPA buffer (Thermo Scientific) supplemented with cOmplete Mini EDTA-free protease inhibitor cocktail (Roche). Post-nuclear lysates were resolved by SDS-PAGE using 4-12% BisTris NuPAGE gels (Invitrogen) and proteins were transferred to membranes (Immunoblot PVDF membranes, Bio-Rad). Membranes were stained with anti-LSP1 (Novus Biologicals) or streptavidin-HRP staining (Pierce). In some experiments, gels were stained with Coomassie Blue (Thermo Fisher).

### Flow cytometry

For the analysis of tonsil cells, tonsil CD14^+^ macrophages were gated as HLA-DR^+^CD11c^+^CD14^+^, tonsil cDC1 as HLA-DR^+^CD11c^+^CD14^−^CD1c^−^CD141^+^ and tonsil cDC2 as HLA-DR^+^CD11c^+^CD14^−^CD1c^+^. For the analysis of blood cells, B cells were gated as HLA-DR^+^CD19^+^, pDC as HLA-DR^+^CD19^−^CD11c^−^CD123^+^, CD16^+^ monocytes as HLA-DR^+^CD19^−^CD16^+^CD14^−^, CD14^+^ monocytes as HLA-DR^+^CD19^−^CD16^−^CD14^+^CD1c^−^, cDC2 as HLA-DR^+^CD19^−^CD16^−^CD14^−^CD1c^+^, DC3 as HLA-DR^+^CD19^−^CD16^−^CD14^int^CD1c^+^.

Antibodies used were: FITC anti-CD19 (clone HIB19, BioLegend), FITC anti-CD3 (clone UCHT1, BioLegend), FITC anti-CD16 (clone 3G8, Biolegend), PE anti-M13 (GE healthcare), PE anti-CD19 (clone SJ25C1, eBioscience), PE anti-CD141 (clone AD5-14H12, Miltenyi biotec), Pe/Cy7 anti-CD11c (clone Bu15, Biolegend), Pe/Cy7 anti-CD163 (clone GHI/61, eBioscience), PerCP-eFluor710 anti-CD1c (clone L161, eBioscience), APC anti-CD304 (clone REA774, Miltenyi biotec), APC anti-CD1a (clone HI149, BioLegend), APC-eFluor780 anti-HLA-DR (clone LN3e, Bioscience), HorizonV500 anti-CD14 (cloneM5E2, BD). In some experiments, cells were stained with Streptavidin-PE (BD), Streptavidin-APC (BD) or Streptavidin-PeCy7 (BD). Fc receptor binding was blocked using TruStain FcX (BioLegend). Dead cells were excluded with 4′,6-diamidino-2-phenylindole (DAPI, Thermo Fisher) staining. Cells were analyzed on a FACSVerse or LSRII (BD Biosciences) instrument. Data was analyzed with FlowJo (Tree Star).

### Immunoprecipitation

All the incubation steps were performed rocking the tubes constantly. Cleared cell lysates resuspended in an equal volume of IP buffer (10 mM Tris–HCl, pH 8.0, 150 mM NaCl, 1% NP40) were pre-incubated 1 h at 4°C in the presence of 200 µL of protein G agarose beads (Thermo Fisher) and successively washed 3 times in IP buffer to eliminate unspecific binding. The supernatant was recovered by centrifugation (3 min × 2500 g), mixed with 200 µg of antibody, and incubated 2 h at 4°C. Finally, 200 µL of washed protein G agarose beads were added and washed after 1 h at 4°C 5 times in 10 mL of IP buffer five times before being resuspended in 50 µL of SDS loading buffer and heated 10 min at 95°C.

### Proteomics Mass-spectrometry

Immunoprecipitation was performed with VHH D4 on *in vitro*-generated mo-DC. Two bands were obtained for the immuno-precipitated material after SDS-PAGE migration (molecular weight around 60 kDa and 45-50 kDa respectively). Gel slices were washed and proteins were reduced with 10 mM DTT before alkylation with 55 mM iodoacetamide. After washing and shrinking the gel pieces with 100% (vol/vol) MeCN, we performed in-gel digestion using trypsin (Roche) overnight in 25 mM NH_4_HCO_3_ at 30 °C. Peptides extracted from each band were analyzed by nanoLC-MS/MS using an Ultimate 3000 system (Dionex, Thermo Scientific, Waltham, MA) coupled to a TripleTOF^TM^ 6600 mass spectrometer (ABSciex). For identification, data was searched against the Swissprot fasta database containing *Homo Sapiens* (2014_10, 20194 sequences) using Mascot^TM^ (version 2.3.02) and further analyzed in myProMS (21). The maximum false discovery rate (FDR) calculation was set to 1% at the peptide level for the whole study (QVALITY algorithm). Only proteins found in two experiments and not in the control IPs were considered candidates. Only one candidate was identified (P33241, Lymphocyte-specific protein 1), with scores of 403.47 and 499.61, and coverage of 23.9% and 26.3%, for the two bands, respectively. Data is available via ProteomeXchange (identifier PXD015647) (22).

## Author contributions

SM, SA, FP and ES designed experiments. SM, AB, AS, BL and ES performed experiments. SM, AB, DL and ES analyzed the data. ES prepared the figures and wrote the manuscript, with input from all authors.

## Acknowledgements

This work was supported by INSERM, CNRS, Agence Nationale de la Recherche (ANR-10-LABX-0043, ANR-CHIN-0002, ANR-10-IDEX-0001-02 PSL), Institut Curie (CIC IGR-Curie 1428), the European Research Council (2013-AdG N° 340046 DCBIOX), Cancéropôle Ile-de-France (2017-1-EMERG-56-ICR-1).

The authors wish to thank the Flow Cytometry Platform of Institut Curie for cell sorting and the TAb-IP platform of Institut Curie for the production of the VHH.

## Notes

**Conflict of interest** SM, SA, FP and ES are co-inventors of a patent entitled "New anti-LSP1 antibody" (PCT/EP2017/0761). The authors declare no other competing interest.

## References

1. Guilliams M, Ginhoux F, Jakubzick C, Naik SH, Onai N, Schraml BU, Segura E, Tussiwand R, and Yona S. Dendritic cells, monocytes and macrophages: a unified nomenclature based on ontogeny. Nat Rev Immunol. 2014;14(8):571–8.

2. Jakubzick CV, Randolph GJ, and Henson PM. Monocyte differentiation and antigen-presenting functions. Nat Rev Immunol. 2017;17(6):349–62.

3. Coillard A, and Segura E. In vivo Differentiation of Human Monocytes. Frontiers in immunology. 2019;10(1907.

4. Ginhoux F, and Guilliams M. Tissue-Resident Macrophage Ontogeny and Homeostasis. Immunity. 2016;44(439–49.

5. Villani AC, Satija R, Reynolds G, Sarkizova S, Shekhar K, Fletcher J, Griesbeck M, Butler A, Zheng S, Lazo S, et al. Single-cell RNA-seq reveals new types of human blood dendritic cells, monocytes, and progenitors. Science. 2017;356(6335).

6. Dutertre CA, Becht E, Irac SE, Khalilnezhad A, Narang V, Khalilnezhad S, Ng PY, van den Hoogen LL, Leong JY, Lee B, et al. Single-Cell Analysis of Human Mononuclear Phagocytes Reveals Subset-Defining Markers and Identifies Circulating Inflammatory Dendritic Cells. Immunity. 2019.

7. Bakdash G, Buschow SI, Gorris MA, Halilovic A, Hato SV, Skold AE, Schreibelt G, Sittig SP, Torensma R, Duiveman-de Boer T, et al. Expansion of a BDCA1+CD14+ Myeloid Cell Population in Melanoma Patients May Attenuate the Efficacy of Dendritic Cell Vaccines. Cancer Res. 2016;76(15):4332–46.

8. Segura E, Touzot M, Bohineust A, Cappuccio A, Chiocchia G, Hosmalin A, Dalod M, Soumelis V, and Amigorena S. Human Inflammatory Dendritic Cells Induce Th17 Cell Differentiation. Immunity. 2013;38(2):336–48.

9. Goudot C, Coillard A, Villani AC, Gueguen P, Cros A, Sarkizova S, Tang-Huau TL, Bohec M, Baulande S, Hacohen N, et al. Aryl Hydrocarbon Receptor Controls Monocyte Differentiation into Dendritic Cells versus Macrophages. Immunity. 2017;47(3):582–96 e6.

10. Moutel S, Bery N, Bernard V, Keller L, Lemesre E, de Marco A, Ligat L, Rain JC, Favre G, Olichon A, et al. NaLi-H1: A universal synthetic library of humanized nanobodies providing highly functional antibodies and intrabodies. Elife. 2016;5(

11. Tang-Huau TL, Gueguen P, Goudot C, Durand M, Bohec M, Baulande S, Pasquier B, Amigorena S, and Segura E. Human in vivo-generated monocyte-derived dendritic cells and macrophages cross-present antigens through a vacuolar pathway. Nature communications. 2018;9(1):2570.

12. Pulford K, Jones M, Banham AH, Haralambieva E, and Mason DY. Lymphocyte-specific protein 1: a specific marker of human leucocytes. Immunology. 1999;96(2):262–71.

13. Durand M, Walter T, Pirnay T, Naessens T, Gueguen P, Goudot C, Lameiras S, Chang Q, Talaei N, Ornatsky O, et al. Human lymphoid organ cDC2 and macrophages play complementary roles in T follicular helper responses. J Exp Med. 2019;216(7):1561–81.

14. Alcantara-Hernandez M, Leylek R, Wagar LE, Engleman EG, Keler T, Marinkovich MP, Davis MM, Nolan GP, and Idoyaga J. High-Dimensional Phenotypic Mapping of Human Dendritic Cells Reveals Interindividual Variation and Tissue Specialization. Immunity. 2017;47(6):1037–50 e6.

15. Klein DP, Jongstra-Bilen J, Ogryzlo K, Chong R, and Jongstra J. Lymphocyte-specific Ca2+-binding protein LSP1 is associated with the cytoplasmic face of the plasma membrane. Mol Cell Biol. 1989;9(7):3043–8.

16. Laoui D, Keirsse J, Morias Y, Van Overmeire E, Geeraerts X, Elkrim Y, Kiss M, Bolli E, Lahmar Q, Sichien D, et al. The tumour microenvironment harbours ontogenically distinct dendritic cell populations with opposing effects on tumour immunity. Nature communications. 2016;7(13720.

17. Lavin Y, Kobayashi S, Leader A, Amir ED, Elefant N, Bigenwald C, Remark R, Sweeney R, Becker CD, Levine JH, et al. Innate Immune Landscape in Early Lung Adenocarcinoma by Paired Single-Cell Analyses. Cell. 2017;169(4):750–65 e17.

18. Michea P, Noel F, Zakine E, Czerwinska U, Sirven P, Abouzid O, Goudot C, Scholer-Dahirel A, Vincent-Salomon A, Reyal F, et al. Adjustment of dendritic cells to the breast-cancer microenvironment is subset specific. Nat Immunol. 2018;19(8):885–97.

19. Jardine L, Wiscombe S, Reynolds G, McDonald D, Fuller A, Green K, Filby A, Forrest I, Ruchaud-Sparagano MH, Scott J, et al. Lipopolysaccharide inhalation recruits monocytes and dendritic cell subsets to the alveolar airspace. Nature communications. 2019;10(1):1999.

20. Durand M, and Segura E. Dendritic Cell Subset Purification from Human Tonsils and Lymph Nodes. Methods Mol Biol. 2016;1423(89–99.

21. Poullet P, Carpentier S, and Barillot E. myProMS, a web server for management and validation of mass spectrometry-based proteomic data. Proteomics. 2007;7(15):2553–6.

22. Vizcaino JA, Csordas A, Del-Toro N, Dianes JA, Griss J, Lavidas I, Mayer G, Perez-Riverol Y, Reisinger F, Ternent T, et al. 2016 update of the PRIDE database and its related tools. Nucleic Acids Res. 2016;44(22):11033.

